# The experimentally obtained functional impact assessments of GT>GC 5’ splice site variants differ markedly from those predicted

**DOI:** 10.1101/864843

**Authors:** Jian-Min Chen, Jin-Huan Lin, Emmanuelle Masson, Zhuan Liao, Claude Férec, David N. Cooper, Matthew Hayden

## Abstract

GT>GC 5’ splice site (or +2T>C) variants have been frequently reported to cause human genetic disease. However, although we have demonstrated that GT>GC variants in human disease genes may not invariably be pathogenic, none of the currently available splicing prediction tools appear to be capable of reliably distinguishing those GT>GC variants that generate wild-type transcripts from those that do not. Recently, SpliceAI, a novel deep residual neural network tool, has been developed for splicing prediction. Methodologically distinct from previous approaches that either rely on human-engineered features and/or which focus on short nucleotide windows adjoining exon-intron boundaries, SpliceAI assesses splicing determinants by evaluating 10,000 nucleotides of flanking contextual sequence to predict the functional role in splicing of each position in the pre-mRNA transcript. Herein, we evaluated the performance of SpliceAI in the context of three datasets of GT>GC variants, all of which had been characterized functionally in terms of their impact on mRNA splicing. The first two datasets refer to our recently described “*in vivo*” dataset of 45 disease-causing GT>GC variants and the “*in vitro*” dataset of 103 GT>GC substitutions. The third dataset comprised 12 *BRCA1* GT>GC variants that were recently analyzed by saturation genome editing. We processed all GT>GC variants using the default settings of SpliceAI. Comparison of the SpliceAI-predicted and experimentally obtained functional impact assessments of the analyzed GT>GC variants revealed that although SpliceAI performed rather better than other prediction tools, it was still far from perfect. A key issue is that the impact of GT>GC (as well as GT>GA or +2T>A) variants that generated wild-type transcripts represents a quantitative change that can vary from barely detectable to almost full expression of wild-type transcripts, with wild-type transcripts often co-existing with aberrantly spliced transcripts. Our findings highlight the challenges that we still face in attempting to accurately identify splice-altering variants.

## 1. INTRODUCTION

Technological advances in DNA sequencing have made whole exome sequencing and even whole genome sequencing increasingly practicable, especially in the clinical setting. However, our ability to accurately interpret the clinical relevance of genetic variants, particularly those that are rare or even private, has so far been quite limited; this represents a rate-limiting step in realizing the full potential of precision medicine (Lappalainen et al., 2019; Shendure et al., 2019). Functional analysis performed in a well-validated assay should provide the strongest possible basis for variant classification (Richards et al., 2015; Starita et al., 2017) but this is often not feasible in practice for certain types of variant. Many computational algorithms have been developed with the aim of predicting the functional effects of different types of genetic variant but none of them meets the exacting standards required in the clinic. This is particularly true for splice-altering variants outside the obligate GT and AG splice-site dinucleotides because (i) splice-altering variants can occur virtually anywhere within a gene’s coding or intronic sequences (Anna and Monika, 2018; Cooper et al., 2009; Scotti and Swanson, 2016; Vaz-Drago et al., 2017) and (ii) splicing is a highly regulated process, involving a complex interaction between *cis*-elements and *trans*-acting factors (Baeza-Centurion et al., 2019; Fu and Ares, 2014; Scotti and Swanson, 2016; Shi, 2017; Wang and Burge, 2008).

Even for variants that occur within the supposedly obligate splice-site dinucleotides, we may still encounter problems of interpretation. For example, variants affecting the 5’ splice site GT dinucleotide, which have been frequently reported to cause human genetic disease (Stenson et al., 2017), are routinely scored as pathogenic splicing mutations and are usually considered to be fully penetrant (Jaganathan et al., 2019; Mount et al., 2019). However, we have recently provided evidence to suggest that 5’ splice site GT>GC variants (henceforth simply termed GT>GC variants or alternatively +2T>C variants) in human disease genes may not invariably be pathogenic (Lin et al., 2019b). Specifically, combining data derived from a meta-analysis of 45 human disease-causing GT>GC variants and a cell culture-based Full-Length Gene Splicing Assay (FLGSA) of 103 GT>GC substitutions, we estimated that ∼15-18% of GT>GC variants generate between 1 and 84% wild-type transcripts (Lin et al., 2019b). During this analysis, we found that none of the four most popular splicing prediction tools, namely SpliceSiteFinder-like, MaxEntScan, NNSPLICE and GeneSplicer (all included within Alamut® Visual; https://www.interactive-biosoftware.com/), were capable of reliably distinguishing those GT>GC variants that generated wild-type transcripts from those that did not (Lin et al., 2019b); for all variants tested, SpliceSiteFinder-like tended to predict a slightly reduced score whilst the other three invariably failed to yield any score. The root of this problem is twofold: Firstly, these splicing prediction tools (in common with many others) focus exclusively on short local DNA sequence motifs and secondly, GC is used instead of GT as the wild-type 5’ splice site dinucleotide in ∼1% of U2 type introns in the human genome (Burset et al., 2000; Parada et al., 2014). It follows that both GT>GC variants that generate wild-type transcripts and those that do not, could in principle occur within identical local sequence tracts as far as the conventional 9-bp 5’ splice site consensus sequence, comprising the last three bases of the preceding exon and the first six bases of the affected intron (the corresponding nucleotide positions are denoted −3_−1/+1_+6), is concerned (Lin et al., 2019b).

Recently, SpliceAI, a novel deep residual neural network tool, has been developed for splicing prediction (Jaganathan et al., 2019). Methodologically distinct from previous approaches that have either relied on human-engineered features and/or focused on short nucleotide windows adjoining exon-intron boundaries, SpliceAI learns splicing determinants directly from the primary sequence by evaluating 10,000 nucleotides of the flanking sequence context to predict the role in splicing of each position in the pre-mRNA transcript. Jaganathan et al. (2019) showed that SpliceAI achieved a top-*k* accuracy of 95% for pre-mRNA transcripts of protein-coding genes and 84% for long intergenic noncoding RNAs (lincRNAs) in the test dataset. [Top-*k* accuracy is defined as the fraction of correctly predicted splice sites at the threshold where the number of predicted sites is equal to the actual number of splice sites present in the test dataset] The accuracy and reliability of SpliceAI was evidenced by (i) the observation that synonymous and intronic variants with predicted splice-altering impact were found to be depleted in the human population, (ii) the fact that 75% of these synonymous and intronic mutations were validated by RNA-seq and (iii) the finding that *de novo* cryptic splice variants were enriched in patients with neurodevelopmental disorders (Jaganathan et al., 2019). Herein, we sought to ascertain whether SpliceAI is capable of accurately distinguishing GT>GC variants that generate wild-type transcripts from those that do not.

## 2. MATERIALS AND METHODS

### 2.1. Source of GT>GC variants

Three datasets of GT>GC variants, all of which have been characterized functionally in terms of their impact on splicing, were employed in this study. The first two datasets correspond to our previously described “*in vivo*” dataset of 45 disease-causing GT>GC variants and the “*in vitro*” dataset of 103 GT>GC substitutions (Lin et al., 2019b). The third dataset comprised 12 GT>GC variants from the *BRCA1* gene, which were extracted from a recent study that prospectively analyzed the functional impact of over 4000 *BRCA1* variants by means of saturation genome editing (Findlay et al., 2018).

In the context of the first dataset (Supplementary Table S1), the precise level of the variant allele-derived wild-type transcripts was available for four of the seven disease-causing GT>GC variants that generated wild-type transcripts in the corresponding original publications (Table 1). For the three remaining variants (i.e., *CAV3* c.114+2T>C in (Muller et al., 2006); *PLP1* c.696+2T>C in (Aoyagi et al., 1999) and *SPINK1* c.194+2T>C in (Kume et al., 2006)), it is apparent from RT-PCR gel photographs in the original publications that all three were associated with the generation of both wild-type and aberrantly spliced transcripts. We employed ImageJ (https://imagej.net) to provide approximate estimates of the levels of the variant allele-derived wild-transcripts for each of the three variants (Table 1).

**Table 1.** Comparison of SpliceAI-predicted and experimentally demonstrated functional effects of the seven disease-causing GT>GC (+2T>C) variants that generated wild-type transcripts

| Gene symbol | mRNA reference | Chromosome | HG38 coordinate | Reference sequence | Variant<sup>a</sup> | % normal expression level<sup>b</sup> | SpliceAI Delta score of donor loss |  |  |
| --- | --- | --- | --- | --- | --- | --- | --- | --- | --- |
|  |  |  |  |  |  |  | +2T>C | +2T>A | +2T>G |
| *CAV3* | NM_001234.4 | 3 | 8733992 | T | c.114+2T>C | 11<sup>c</sup> | 0.9 | 1 | 1 |
| *CD3E* | NM_000733.3 | 11 | 118313876 | T | c.520+2T>C | 1-5<sup>d</sup> | 0.99 | 0.99 | 0.99 |
| *CD40LG* | NM_000074.2 | X | 136654432 | T | c.346+2T>C | 15<sup>d</sup> | 0.95 | 0.97 | 0.97 |
| *DMD* | NM_004006.2 | X | 31657988 | A | c.8027+2T>C | 10<sup>d</sup> | 0.63 | 0.99 | 0.99 |
| *PLP1* | NM_000533.4 | X | 103788512 | T | c.696+2T>C | 8<sup>c</sup> | 0.74 | 1 | 1 |
| *SLC26A2* | NM_000112.3 | 5 | 149960981 | T | c.-26+2T>C | 5<sup>d</sup> | 0.9 | 0.99 | 0.99 |
| *SPINK1* | NM_003122.3 | 5 | 147828020 | A | c.194+2T>C<sup>e</sup> | 10<sup>c</sup> | 0.35 | 0.99 | 1 |

<sup>a</sup>Nomenclature in accordance with Human Genome Variation Society (HGVS) recommendations (den Dunnen et al., 2016).
<sup>b</sup>Expressed as the level of the variant allele-derived wild-type transcripts relative to that of the wild-type allele-derived wild-type transcripts.
<sup>c</sup>Expression level determined here by ImageJ using gel photos from the original publications.
<sup>d</sup>Expression level as described in the original publications.
<sup>e</sup>Identical to the *SPINK1* IVS3+2T>C substitution in Table 2.

### 2.2. Variant description and nomenclature

Variant description and nomenclature were in line with our previous publication (Lin et al., 2019b). First, we used the term ‘variants’ to describe naturally occurring disease-causing events and ‘substitutions’ to denote artificially engineered events. Second, 5’ splice site GT>GC, GT>GA and GT>GG variants or substitutions were used synonymously with +2T>C, +2T>A and +2T>G variants or substitutions, respectively. Third, disease-causing variants were named in accordance with Human Genome Variation Society (HGVS) recommendations (den Dunnen et al., 2016) whilst the traditional IVS (InterVening Sequence; i.e., an intron) nomenclature was used for the engineered substitutions. Finally, hg38 positions (https://genome.ucsc.edu/) for all variants or substitutions under study are systematically provided in the various tables.

### 2.3. SpliceAI prediction

GT>GC variants or substitutions as well as their corresponding GT>GA and GT>GG counterparts were processed (during October 2019) using the default settings of SpliceAI version 1.2.1, with a custom gene annotation file containing NCBI reference sequence transcript start and end coordinates. Default settings, and instruction for use of custom annotation files, were taken from https://pypi.org/project/spliceai/.

### 2.4. Performance testing

Two statistical tests, a Matthews correlation coefficient (MCC) and a Receiver operating characteristic (ROC) curve, were carried out on the dataset 2 substitutions assessed by SpliceAI. MCC test is a correlation coefficient between the observed and predicted binary classifications. For a perfect prediction, the coefficient is +1; a coefficient of 0 is no better than random, and no correlation between observed and predicted yields −1 (Matthews, 1975). A ROC curve illustrates the diagnostic specificity and sensitivity of a binary classifier system as its discrimination threshold is varied; this enables the selection of an optimum threshold value. To assess the difference between the diagonal and the ROC curve obtained, the area under the ROC curve is measured (AUC). An AUC of 0.5 would be a random prediction whilst a perfect predictor would score 1. ROC analysis was carried out using the R-based web tool easyROC (Goksuluk et al., 2016).

For the MCC test, a contingency table was derived from dataset 2 (Supplementary Table S2) where a true positive is defined as a predicted splice altering substitution for which FLGSA produced no transcript and a true negative is a substitution not predicted to alter splicing and for which FLGSA produces transcript.

### 2.5. Functional analysis of two GT-affecting variants

The functional impact of two newly engineered GT-affecting variants in the *HESX1* gene were analyzed by means of the cell culture-based FLGSA method as previously described (Lin et al., 2019b).

## 3. RESULTS AND DISCUSSION

### 3.1. Accuracy and reliability of the experimentally obtained functional assessment of the GC>GT variants analyzed

We employed SpliceAI to make predictions as to the splicing consequences of GT>GC variants from three distinct datasets. Since the experimentally ascertained functional impact of the GT>GC variants analyzed was used as the starting point for our analysis, their accuracy and reliability were of critical importance. Regarding the first dataset of known pathogenic variants (Supplementary Table S1), several points are worth highlighting. First, all 45 disease-causing variants were either homozygotes, hemizygotes or compound heterozygotes, a prerequisite for determining the presence or absence of the variant allele-derived normal transcripts. Second, for each variant, patient-derived tissue or cells (pathologically relevant in about half of the cases) had been used to perform the RT-PCR analysis that had unequivocally demonstrated the presence or absence of variant allele-derived wild-type transcripts in the corresponding original publication. Third, the levels of the variant allele-derived wild-type transcripts in the seven disease-causing GT>GC variants that generated wild-type transcripts were very low (≤15% of normal; Table 1), potentially explicable by the ascertainment bias inherent to all disease-causing variants. Nonetheless, all seven of these variants were noted to be associated with a milder clinical phenotype than would have been expected from a functionally null variant (Lin et al., 2019b), consistent with other findings that even the retention of a small fraction of normal gene function can significantly impact the clinical phenotype (Den Uijl et al., 2011; Ramalho et al., 2002; Raraigh et al., 2018; Scalet et al., 2019).

In the case of the second dataset (Supplementary Table S2), the functional effects of all 103 engineered GT>GC substitutions (from 30 different genes) were analyzed by Full-Length Gene Splicing Assay (FLGSA) in transfected HEK293T cells (Lin et al., 2019b), with all 19 substitutions that generated some wild-type transcripts being listed in Table 2. By comparison to the commonly used minigene splicing assay, FLGSA preserves better the natural genomic sequence context of the studied variants (Wu et al., 2017; Zou et al., 2016). The accuracy and reliability of the FLGSA-derived data can be inferred from the following three lines of evidence. First, 10 GT>GC substitutions that generated wild-type transcripts and 10 GT>GC substitutions that did not generate wild-type transcripts in transfected HEK293T cells were further analyzed in transfected HeLa cells using FLGSA, yielding entirely consistent findings in terms of whether or not wild-type transcripts were generated (Lin et al., 2019b). Second, *HESX1* c.357+2T>C and *SPINK1* c.194+2T>C were the only variants common to both the first and second datasets. The functional effects of these two variants *in vivo* — *HESX1* c.357+2T>C generated no wild-type transcripts whereas *SPINK1* c.194+2T>C generated some wild-type transcripts (Supplementary Table S1) — were faithfully replicated in FLGSA (Supplementary Table S2). Third, a GT>GC variant that was not present in either dataset, *HBB* c.315+2T>C, had been reported to be associated with a milder hematological phenotype and it was suggested that it might have a limited impact on splicing (Frischknecht et al., 2009). Using FLGSA performed in HEK293T cells, we found that it generated a low level of wild-type transcripts (Lin et al., 2019b). Importantly, the orthologous variant of *HBB* c.315+2T>C in the rabbit *Hbb* gene has also been found to be capable of generating wild-type transcripts in two experimental model systems, namely *in vitro* transcription of *Hbb* RNA in a HeLa cell nuclear extract and transient expression of the full-length *Hbb* gene in HeLa cells (Aebi et al., 1986; Aebi et al., 1987). These notwithstanding, tissue-or cell-specific factors have on some occasions impacted splicing (Jaganathan et al., 2019; Pineda and Bradley, 2018), an issue that was not extensively addressed in our previous study (Lin et al., 2019b). The bottom line here is that (i) the 30 genes used for FLGSA analysis were selected using a procedure that did not take into consideration the gene’s function or expression, (ii) all 30 genes underwent normal splicing in the context of their reference mRNA sequences as specified in Supplementary Table S2 and(iii) the generation (or not) of wild-type transcripts from the engineered GC allele was observed under the same experimental conditions as for the wild-type GT allele (Lin et al., 2019b).

**Table 2.** Comparison of SpliceAI-predicted and experimentally demonstrated functional effects of the 19 engineered GT>GC (+2T>C) substitutions that generated wild-type transcripts

| Gene symbol | mRNA reference | Chromosome | hg38 coordinate | Reference sequence | Substitution<sup>a</sup> | Generation of wild-type transcripts<sup>b</sup> | SpliceAI Delta score of donor loss |  |  |
| --- | --- | --- | --- | --- | --- | --- | --- | --- | --- |
|  |  |  |  |  |  |  | +2T>C | +2T>A | +2T>G |
| *CCDC103* | NM_213607.2 | 17 | 44899861 | T | IVS1+2T>C | Yes | 0.82 | 0.82 | 0.82 |
| *DBI* | NM_001079862.2 | 2 | 119368307 | T | IVS2+2T>C | Yes | 0.86 | 1 | 1 |
| *DNAJC19* | NM_145261.3 | 3 | 180985924 | A | IVS5+2T>C | Yes (42%) | 0.03 | 0.99 | 0.95 |
| *FATE1* | NM_033085.2 | X | 151716227 | T | IVS1+2T>C | Yes (84%) | 0.08 | 0.96 | 1 |
| *FOLR3* | NM_000804.3 | 11 | 72139484 | T | IVS4+2T>C | Yes | 0.45 | 1 | 1 |
| *HESX1* | NM_003865.2 | 3 | 57199760 | A | IVS1+2T>C | Yes (2%) | 0.81 | 0.98 | 0.98 |
| *IFNL2* | NM_172138.1 | 19 | 39269823 | T | IVS5+2T>C | Yes (5%) | 0.05 | 0.84 | 0.73 |
| *IL10* | NM_000572.3 | 1 | 206770905 | A | IVS3+2T>C | Yes | 0.61 | 1 | 1 |
| *MGP* | NM_000900.4 | 12 | 14884211 | A | IVS2+2T>C | Yes (80%) | 0.97 | 0.99 | 0.99 |
| *PSMC5* | NM_001199163.1 | 17 | 63830503 | T | IVS6+2T>C | Yes (56%) | 0.31 | 0.98 | 1 |
|  |  |  | 63831228 | T | IVS8+2T>C | Yes (56%) | 0.21 | 1 | 1 |
|  |  |  | 63831618 | T | IVS10+2T>C | Yes (46%) | 0.83 | 1 | 1 |
| *RPL11* | NM_000975.5 | 1 | 23692761 | T | IVS2+2T>C | Yes | 0 | 0.87 | 0.86 |
|  |  |  | 23693915 | T | IVS3+2T>C | Yes | 0.74 | 1 | 1 |
| *RPS27* | NM_001030.4 | 1 | 153991225 | T | IVS2+2T>C | Yes (63%) | 0.67 | 1 | 1 |
|  |  |  | 153991678 | T | IVS3+2T>C | Yes | 0.98 | 1 | 1 |
| *SELENOS* | NM_203472.2 | 15 | 101277340 | A | IVS1+2T>C | Yes | 0.81 | 1 | 1 |
|  |  |  | 101274418 | A | IVS5+2T>C | Yes (14%) | 0.79 | 1 | 1 |
| *SPINK1* | NM_003122.3 | 5 | 147828020 | A | IVS3+2T>C<sup>c</sup> | Yes | 0.35 | 0.99 | 1 |

<sup>a</sup>In accordance with the traditional IVS (InterVening Sequence; i.e., an intron) nomenclature as previously described (Lin et al. 2019b).
<sup>b</sup>Expression level (in parentheses), determined by quantitative RT-PCR analysis, was available for all +2T>C substitutions that generated only wild-type transcripts under the experimental conditions described in (Lin et al. 2019b).
<sup>c</sup>Identical to the *SPINK1* c.194+2T>C variant in Supplementary Table S1 and Table 1.

The third dataset was obtained courtesy of a perusal of the literature (Table 3). Recently, the functional impact of all possible single nucleotide substitutions within 13 exons and adjacent intronic sequences of the 23-exon *BRCA1* gene (NM_007294.3) have been prospectively analyzed by means of saturation genome editing (Findlay et al., 2018). Taking advantage of the essentiality of *BRCA1* in the human near-haploid cell line HAP1 (Blomen et al., 2015), Findlay and colleagues used cell viability as a proxy indicator for the functional consequences of the analyzed substitutions. It should be noted that the functional consequences of all tested substitutions were actually evaluated in their natural genomic sequence contexts. Of the ∼4000 *BRCA1* single nucleotide substitutions analyzed, 12 were GT>GC substitutions. Of these 12 GT>GC substitutions, one was classified as “functional”, two were classified as “intermediate” and the remaining nine were classified as “non-functional” (Table 3). Whereas “functional” and “intermediate” were interpreted as having generated wild-type transcripts, “non-functional” was interpreted as having not generated any wild-type transcripts (Lin et al., 2019a). As such, 25% (n = 3) of these 12 *BRCA1* GT>GC substitutions generated wild-type transcripts, a proportion largely consistent with our estimated 15-18% rate. Moreover, the *BRCA1* GT>GC variant in intron 18 was shown to be “functional”, providing further support for our contention that GT>GC variants in human disease genes may not invariably be pathogenic (Lin et al., 2019b).

**Table 3.**
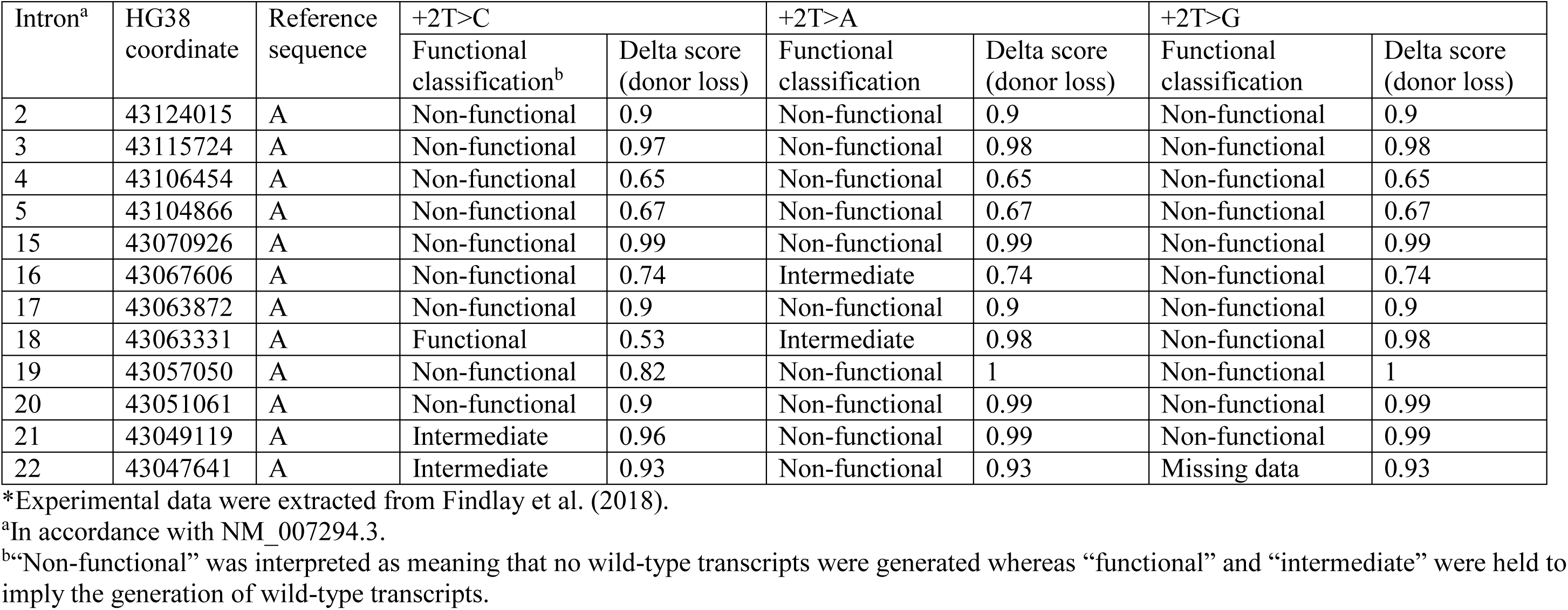
Comparison of SpliceAI-predicted and experimentally demonstrated functional effects of all possible single nucleotide substitutions in the +2 positions of 12 *BRCA1* introns*

Taken together, the experimentally obtained functional assessments of the included GC>GT variants or substitutions were considered to be of high quality and appropriate for the intended study.

### 3.2. Selection and interpretation of SpliceAI Delta scores for analysis

We processed GT>GC variants using the default settings of SpliceAI as detailed in https://pypi.org/project/spliceai/. SpliceAI provides Delta scores (ranging from 0 to 1) for each variant, thereby providing a measure of their probability of altering splicing in terms of either splice donor gain, splice donor loss, splice acceptor gain, and splice acceptor loss. SpliceAI also provides Delta position that conveys information specifying the location where splicing differs from normal relative to the position of the associated variant. Since the GT>GC variants or substitutions under study invariably affected the +2 position of the canonical 5’ splice site GT dinucleotides (in the context of the specified mRNA reference sequence), we focused our analysis exclusively on the Delta scores of donor loss although other scores may provide clues as to the nature of the resulting aberrantly spliced transcripts of splice-altering variants. Thus, only the SpliceAI-predicted Delta scores of donor loss for the studied GT>GC variants or substitutions are provided in Supplementary Tables S1 and S2 as well as in Tables 1-3. Here, it is important to note two points. First, the previously studied GT>GC events generated maximally 84% wild-type transcripts as compared to their wild-type GT allele counterparts (Lin et al., 2019b). In other words, all these variants were associated minimally with a 16% functional loss. Therefore, strictly speaking, all these previously studied GT>GC events can be defined as splice-altering. Second, in those cases of GT>GC events that generated wild-type transcripts, the level of wild-type transcript varied from 1-84% (Lin et al., 2019b). Intuitively, whether or not a GT>GC variant capable of generating wild-type transcripts is pathogenic is likely to depend at least in part upon the level of the generated wild-type transcripts. Taking these points into consideration, we shall use the SpliceAI Delta score of donor loss as a proxy indicator of the probability of a given GT>GC variant being able to generate wild-type transcripts; variants with a Delta score above a certain cutoff value will be considered not to be capable of generating wild-type transcripts whereas variants with a Delta score below the cutoff value will be considered as being capable of generating wild-type transcripts.

### 3.3. Encouraging findings from a quick survey of the three datasets of GT>GC variants

As mentioned in the Introduction, none of the four most popular splicing prediction tools, SpliceSiteFinder-like, MaxEntScan, NNSPLICE and GeneSplicer, were found to be able to distinguish those GT>GC variants that generated wild-type transcripts from those that did not (Lin et al., 2019b). As described below, a quick survey of SpliceAI-predicted scores yielded encouraging results across all three datasets of GT>GC variants.

First, in the context of dataset 1, the level of variant allele-derived wild-type transcripts associated with the seven disease-causing GT>GC variants was at most 15% of normal (Table 1). Although this low level increase in the generation of wild-type transcripts may make prediction a daunting task, it is interesting to see that the two lowest Delta scores of donor loss, 0.35 and 0.63, were observed in association with the two variants that generated ∼10% wild-type transcripts (Supplementary Table S1; Table 1). The score of 0.35 was observed for the *SPINK1* c.194+2T>C variant, for which the RT-PCR analysis was performed using gastric tissue from a homozygous patient with chronic pancreatitis (Kume et al., 2006). Although stomach is not known to be clinically affected in chronic pancreatitis, the expression data were considered to be highly reliable for two reasons. Firstly, the *in vivo* expression data was confirmed by FLGSA performed in both HEK293T and HeLa cells (Lin et al., 2019b). Secondly, had the *SPINK1* c.194+2T>C variant in question caused a complete functional loss of the affected allele, the homozygotes should have developed severe infantile isolated exocrine pancreatic insufficiency instead of chronic pancreatitis (Venet et al., 2017). The score of 0.63 was observed for the *DMD* c.8027+2T>C variant, for which the detection of wild-type transcripts was based upon RT-PCR analysis of disease-affected muscle tissue from a hemizygous carrier with Becker muscular dystrophy (Bartolo et al., 1996).

As for the second dataset (Supplementary Table S2), the four lowest Delta scores of donor loss (i.e., 0, 0.03, 0.05 and 0.08) were all found in substitutions that generated wild-type transcripts; and 63% (n = 12) of the 19 substitutions that generated some wild-type transcripts had a Delta score of <0.80 (Table 2). As for the third dataset, the lowest Delta score, 0.53, was observed in association with the only “functional” *BRCA1* IVS18+2T>C variant (Table 3).

### 3.4. Statistical comparison of experimentally obtained functional data with SpliceAI predictions for the 103 engineered GT>GC splice variants (dataset 2)

Dataset 2 comprised 19 substitutions that generated wild-type transcripts and 84 substitutions that generated no wild-type transcripts. We thus performed two statistical tests, a Receiver operating characteristic (ROC) curve and a Matthews correlation coefficient (MCC), on the 103 substitutions assessed by SpliceAI (Supplementary Table S2) with a view both to identifying an optimum threshold value and to assessing the correlation between the FLGSA assay results and the SpliceAI predictions.

Based on an ROC analysis of 103 variants from dataset 2 (Supplementary Table S2), an optimum threshold point of 0.85 was provided - similar to the threshold of 0.80, recommended by SpliceAI for high precision results. A contingency table was constructed (Supplementary Table S3) to calculate values for the false positive rate, specificity, sensitivity, accuracy and the Matthews correlation coefficient. These are summarized in Table 4, along with the AUC result obtained from the ROC analysis, the curve from which is shown in Fig. 1.

**Table 4.**
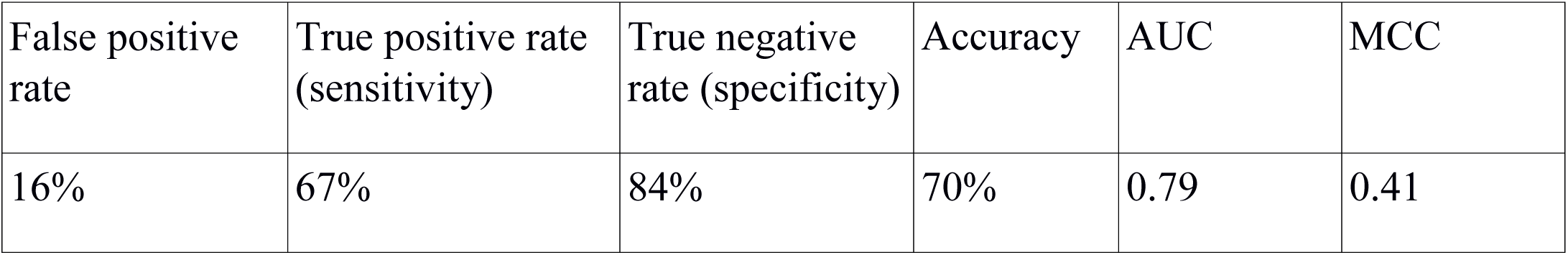
Performance metrics of SpliceAI as a predictor for splice site disruption on 103 variants from dataset 2

| False positive rate | True positive rate (sensitivity) | True negative rate (specificity) | Accuracy | AUC | MCC |
| --- | --- | --- | --- | --- | --- |
| 16% | 67% | 84% | 70% | 0.79 | 0.41 |

**Figure 1.**
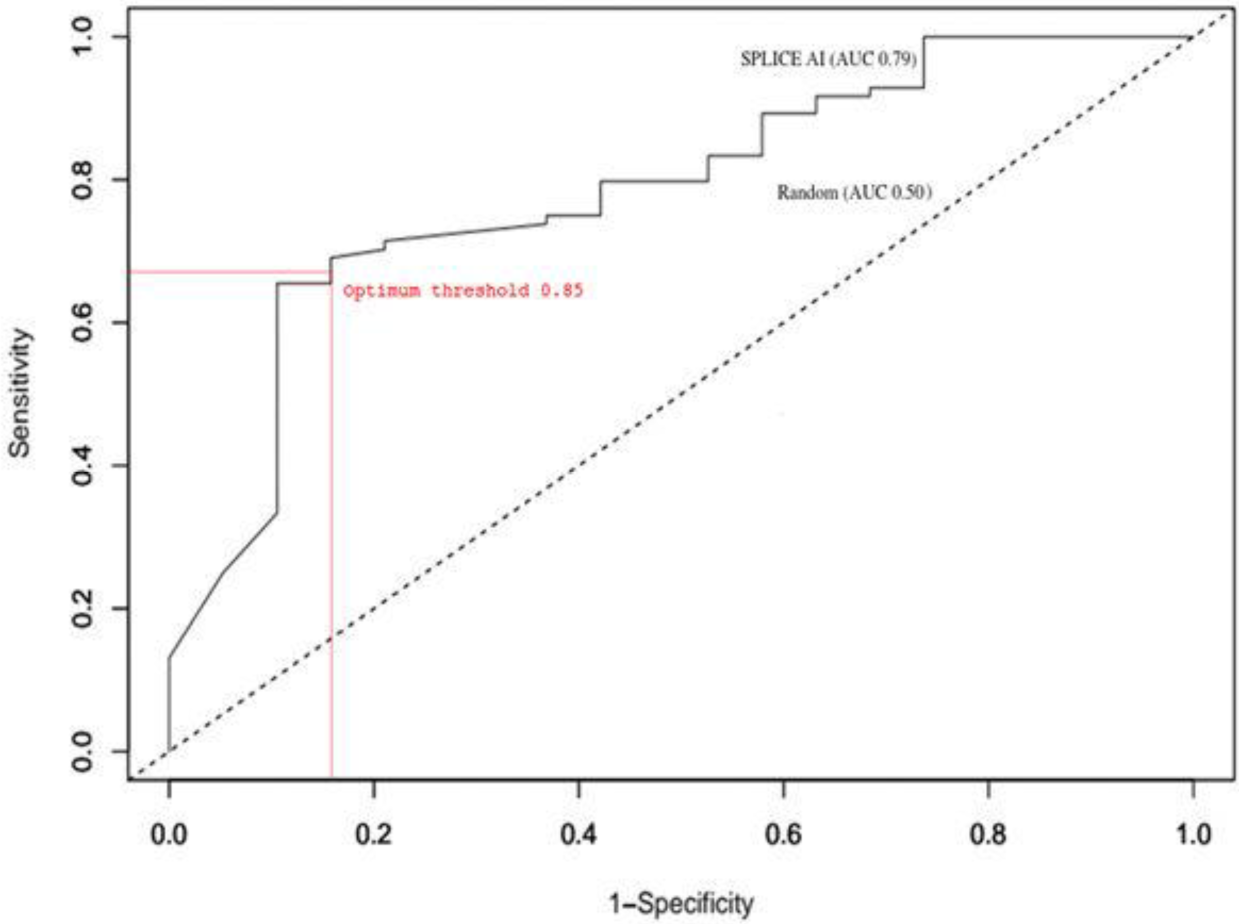
A receiver operating characteristic (ROC) curve for the SpliceAI predictions generated from dataset 2 (Supplementary Table S2), with dotted diagonal line indicating a random prediction (0.5 AUC) and the solid line showing SpliceAI prediction performance (0.79 AUC). The intersection between the two represents the optimum threshold.

As can be seen from Table 4, the AUC of 0.79 and the MCC score of 0.41 are indicative of a good correlation between predicted and actual results. There is also a low false positive rate whilst still maintaining a high accuracy and sensitivity. These results show that for dataset 2, at a threshold of 0.85, SpliceAI can accurately discriminate between those GT>GC substitutions which disrupt splicing and transcript production and those which do not disrupt splicing and produce transcript.

### 3.5. Considerable discrepancy between the predicted and experimentally obtained functional impact assessments of GT>GC 5’ splice site variants

Employing 0.85 as the threshold Delta score (donor loss) to define the generation of wild-type transcripts, rather variable performance between SpliceAI-predicted and experimentally demonstrated functional effects of GT>GC variants were observed across the three datasets: 33-84% of the variants that generated wild-type transcripts and 67-89% of the variants that generated no wild-type transcripts were correctly predicted by SpliceAI (Table 5).

**Table 5.** Overall correlation rates between SpliceAI-predicted and experimentally demonstrated functional effects of the GT>GC variants in the context of three datasets*

| **Variants generating wild-type transcripts** |  |
| --- | --- |
| Dataset 1 (45 disease-causing variants) | 43% (3/7) |
| Dataset 2 (103 variants analyzed by FLGSA) | 84% (16/19) |
| Dataset 3 (12 *BRCA1* variants analyzed by saturation genome editing) | 33% (1/3) |
| **Variants generating no wild-type transcripts** |  |
| Dataset 1 (45 disease-causing variants) | 89% (34/38) |
| Dataset 2 (103 variants analyzed by FLGSA) | 68% (57/84) |
| Dataset 3 (12 *BRCA1* variants analyzed by saturation genome editing) | 67% (6/9) |

\*Splice AI Delta score (donor loss) of 0.85 was used as the threshold value for defining the generation of wild-type transcripts or not.

The poorest performance (43% (3/7) and 33% (1/3)) was observed with datasets 1 and 3 variants that generated wild-type transcripts (Table 5). In the context of the seven dataset 1 variants that generated wild-type transcripts (a qualitative property), the relatively poor performance of 43% might be related to the fact that the functional impact of these GT>GC variants actually manifested as rather small quantitative changes, generating between 1-15% normal transcripts (Table 1). This notwithstanding, it should be pointed out that the two disease-causing variants that generated 10-15% wild-type transcripts, *CAV3* c.114+2T>C (Muller et al., 2006) and *CD40LG* c.346+2T>C (Seyama et al., 1998), had Delta scores of ≥0.9 (Table 1); and in each of these two cases, RT-PCR analysis was performed using patient-derived and pathologically relevant tissue or cells. In short, it remains unclear why some of the disease-causing variants that generated comparable levels of wild-type transcripts were predicted to have low Delta scores (i.e., *DMD* c.8027+2T>C and *SPINK1* c.194+2T>C) whereas others were predicted to have high Delta scores (i.e., *CAV3* c.114+2T>C and *CD40LG* c.346+2T>C). In the context of dataset 3 substitutions that generated wild-type transcripts, the precise levels of wild-type transcripts generated by the two “intermediate” *BRCA1* +2T>C substitutions (both had a Delta score of ≥0.93; Table 3) were unknown.

As for variants that did not generate wild-type transcripts, an excellent correlation rate, 89%, was observed with the 38 such disease-causing variants. By contrast, the performance in datasets 2 and 3 variants was much lower and almost identical (68% and 67%, respectively; Table 5). A fundamental difference between dataset 1 variants and the latter two dataset substitutions is that all of the former were previously published whilst almost all of the latter were prospectively generated. Thus, it is tempting to speculate that for most of the 38 disease-causing variants that did not generate wild-type transcripts, their functional effects might have been ‘seen’ by SpliceAI during training, thereby leading to a higher correlation rate.

In an attempt to further understand the poor performance of dataset 2 and 3 substitutions that did not generate wild-type transcripts, we opted to use the corresponding +2T>A and +2T>G substitutions as controls. The underlying premise was that, based upon current knowledge, +2T>A and +2T>G variants should completely disrupt normal splicing and consequently have high Delta scores in virtually all cases (see also section 3.6). Here it is worth mentioning that we previously employed FLGSA to analyze the functional impact of 15 +2T>A substitutions and 18 +2T>G substitutions, none of which generated any wild-type transcripts (Lin et al., 2019b). We processed all corresponding +2T>A and +2T>G variants by means of SpliceAI in the same way as for the +2T>C variants (during October 2019), the resulting Delta scores for donor loss being provided in Tables 1-3 and Supplementary Tables S1 and S2.

As shown in Supplementary Table S1, all +2T>A and +2T>G variants corresponding to the 45 disease-causing +2T>C variants had very high Delta scores, ranging from 0.92 to 1. By contrast, 91% (n = 94) of the +2T>A and +2T>G variants corresponding to the 103 dataset 2 +2T>C substitutions had a Delta score of ≥0.85 (Supplementary Table S2). In other words, nine of the 103 +2T sites were predicted to have a Delta score of <0.85 when substituted by either A or G; and in these sites, the Delta scores are often identical for all three possible substitutions (Table 6). One possible reason for lower than expected Delta scores is provided in (Jaganathan et al., 2019); exons which undergo a substantial degree of alternative splicing, defined as being between 10% and 90% exon inclusion averaged across samples, tend towards intermediate scores (stated as between 0.35 and 0.8). We therefore explored this possibility using the two sites for which all possible substitutions had the lowest Delta scores (i.e., 0.59 and 0.3; Table 6) as examples. To this end, alternative transcripts of the genes of interest were surveyed via https://www.ncbi.nlm.nih.gov/gene/.

**Table 6.** Nine +2T positions for which all three possible nucleotide substitutions had a consistent SpliceAI Delta score of <0.85

| Gene symbol | mRNA reference | Chromosome | hg38 coordinate | Reference sequence | SpliceAI Delta score of donor loss |  |  |
| --- | --- | --- | --- | --- | --- | --- | --- |
|  |  |  |  |  | +2T>C | +2T>A | +2T>G |
| *AURKC* | NM_001015878.1 | 19 | 57235060 | T | 0.8 | 0.8 | 0.8 |
| *CCDC103* | NM_213607.2 | 17 | 44899861 | T | 0.82 | 0.82 | 0.82 |
| *FABP7* | NM_001446.4 | 6 | 122779869 | T | 0.83 | 0.84 | 0.84 |
| *IFNL2* | NM_172138.1 | 19 | 39269823 | T | 0.05 | 0.84 | 0.73 |
| *LY6G6F* | NM_001003693.1 | 6 | 31708136 | T | 0.81 | 0.81 | 0.81 |
|  |  |  | 31710420 | T | 0.3 | 0.3 | 0.3 |
| *PSMC5* | NM_001199163.1 | 17 | 63830191 | T | 0.76 | 0.77 | 0.77 |
| *RPL11* | NM_000975.5 | 1 | 23695910 | T | 0.59 | 0.59 | 0.59 |
| *SELENOS* | NM_203472.2 | 15 | 101272762 | A | 0.64 | 0.64 | 0.64 |

All three possible single nucleotide substitutions in the *RPL11* g.23695910T (IVS5+2T in accordance with NM_000975.5) site had an identical Delta score of 0.59. *RPL11* has two transcripts, the other being NM_001199802.1. Nonetheless, the two transcripts have common coding sequences from exons 3-6. Moreover, all three possible single nucleotide substitutions in the NM_000975.5-defined *RPL11* IVS5+2T site have been previously subjected to FLGSA, invariably generating no wild-type transcripts (Lin et al., 2019b). Taken together, in this particular case, the lower than expected Delta scores cannot be adequately explained by alternative splicing.

All three possible single nucleotide substitutions in the *LY6G6F* g.31708136T (IVS5+2T in accordance with NM_001003693.1) site had a score of 0.3 (Table 6). NM_001003693.1-defined *LY6G6F* has sequence from exons 1 to 4 in common with NM_001353334.2-defined *LY6G6F-LY6G6D*, which represents naturally occurring readthrough transcription between the neighboring *LY6G6F* and *LY6G6D* genes on chromosome 6 (Supplementary Fig. S1). By contrast, NM_001003693.1-defined exons 5 and 6 are spliced out in NM_001353334.2-defined *LY6G6F-LY6G6D*. It is likely that the use of the “*LY6G6F* IVS5+2T site” as a splice site in one transcript isoform but not in the other underlies the similarly low Delta scores for the three above mentioned possible single nucleotide substitutions. However, two points should be emphasized here. Firstly, none of the three possible single nucleotide substitutions in the context of the NM_001003693.1-defined *LY6G6F* IVS5+2T site led to the generation of wild-type transcripts as evidenced by FLGSA. Whether these substitutions would lead to the increased use of NM_001353334.2-defined *LY6G6F-LY6G6D* remains unclear. Secondly, all three possible single nucleotide substitutions, if considered only in the context of NM_001353334.2-defined *LY6G6F-LY6G6D* (Supplementary Fig. S1), may not affect gene splicing at all.

Finally, let us turn our attention to the *BRCA1* findings in relation to NM_007294.3 (Table 3). The lowest Delta score of donor loss in the context of +2T>A and +2T>G variants, 0.65, was observed for all three possible SNVs in the *BRCA1* IVS4+2T site. The next lowest score, 0.67, was observed for all three possible SNVs in the *BRCA1* IVS5+2T site (Table 3). All six of these variants have been analyzed using saturation genome editing and were invariably classified as “non-functional” (Jaganathan et al., 2019). Moreover, although *BRCA1* has multiple transcripts, these two introns are used by all transcripts (Supplementary Fig. S2). Therefore, as in the abovementioned *RPL11* case, these lower than expected Delta scores cannot be adequately explained by alternative splicing.

### 3.6. Additional findings

We succeeded in analyzing two additional engineered GT-impacting substitutions in the *HESX1* gene, IVS2+2T>A (hg38# chr3:57198751A>T) and IVS3+2T>G (hg38# chr3:57198389A>C), using the cell culture-based FLGSA method. Interestingly, the IVS2+2T>A substitution generated both wild-type and aberrant transcripts whereas IVS3+2T>G generated only aberrant transcripts (Fig. 2). Moreover, two of the 12 *BRCA1* +2T>A substitutions, IVS16+2T>A and IVS18+2T>A, were described as being “intermediate” (Table 3). Although no disease-causing +2T>A variants have been found to generate wild-type transcripts, GA has recently been ranked fourth in terms of its relative frequency among the six noncanonical 5’ splice sites identified by genome-wide RNA-seq analysis and splicing reporter assays (Erkelenz et al., 2018). However, of the three +2T>A substitutions that were experimentally shown to generate some wild-type transcripts, two were predicted to have a Delta score of >0.85, namely 0.93 for *HESX1* IVS2+2T>A (Supplementary Table S2) and 0.98 for *BRCA1* IVS18+2T>A (Table 3). The other one, *BRCA1* IVS16+2T>A, was predicted to have a Delta score of 0.74; but an identical score was also predicted for *BRCA1* IVS16+2T>C and IVS16+2T>G, both of which were classified as “non-functional” (Table 3). In short, SpliceAI appeared not to work as well for the +2T>A variants that generated wild-type transcripts as for the +2T>C variants that generated wild-type transcripts.

**Figure 2.**
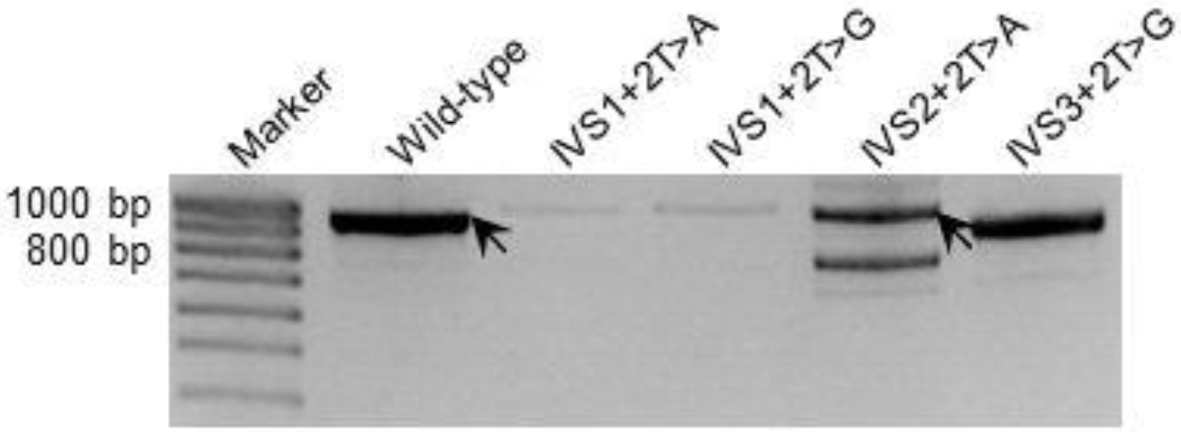
RT-PCR analyses of HEK293T cells transfected with full-length *HESX1* gene expression constructs carrying respectively the wild-type and four different nucleotide substitutions. Wild-type transcripts emanating from the wild-type construct and the construct containing the IVS2+2T>A substitution (confirmed by DNA sequencing) are indicated by arrows. IVS2+2T>A (hg38# chr3:57198751A>T) and IVS3+2T>G (hg38# chr3:57198389A>C) substitutions were newly analyzed here. IVS1+2T>A and IVS1+2T>G, which had been previously analyzed (Lin et al., 2019b), are included for the sake of comparison.

## 4. CONCLUSIONS AND PERSPECTIVES

In the present study, we attempted to correlate SpliceAI-predicted and experimentally obtained functional effects of GT>GC variants in the context of three independent and complementary datasets. Employing data from dataset 2 substitutions, we were able to propose a Delta score of donor loss, 0.85, as defining the threshold of whether or not wild-type transcripts would be generated by GT>GC variants; whereas a score of ≥0.85 defines the complete absence of wild-type transcripts, a score of <0.85 defines the generation of at least some wild-type transcripts. Subsequent use of this threshold score to correlate SpliceAI-predicted and experimentally obtained functional effects of the GT>GC variants revealed that SpliceAI performed better than the popular prediction tools such as SpliceSiteFinder-like, MaxEntScan, NNSPLICE and GeneSplicer. However, a considerable discrepancy still existed between SpliceAI-predicted and experimentally obtained functional assessments in relation to GT>GC (as well as GT>GA) variants. Indeed, this discrepancy serves to illuminate the challenges ahead in accurately identifying all splice-altering variants. A key issue here is that the impact of GT>GC (as well as GT>GA) variants that generated wild-type transcripts represents a quantitative change that can vary from barely detectable to almost full expression of wild-type transcripts, with wild-type transcripts often co-existing with aberrantly spliced transcripts. This is also the case for most of the splice-altering variants occurring outside the essential splice site dinucleotides, whose effects “are not fully penetrant and a mixture of both normal and aberrant splice isoforms are produced” (Jaganathan et al., 2019). Moreover, there is also the issue of alternative splicing related to tissue- or cell-specific factors. While it is clear that we are still very far acquiring a full understanding of the ‘splicing code’ (Bao et al., 2019), we are of the opinion that any improvement in the prioritization of splicing variants will necessitate the refinement of *in silico* prediction tools by reference to *in vitro* functional assessment courtesy of the results obtained from well-validated assays such as FLGSA.

## ACKNOWLEDGMENTS

We are grateful to the original authors who reported the disease-causing 5’ splice site GT>GC variants studied here. J.H.L. was in receipt of a 20-month scholarship from the China Scholarship Council (No. 201706580018). This study was supported by the Institut National de la Santé et de la Recherche Médicale (INSERM), France. D.N.C. and M.H. acknowledge the financial support of Qiagen plc through a License Agreement with Cardiff University.

## REFERENCES

Aebi M, Hornig H, Padgett RA, Reiser J, Weissmann C. Sequence requirements for splicing of higher eukaryotic nuclear pre-mRNA. Cell 1986; 47: 555–565.

Aebi M, Hornig H, Weissmann C. 5’ cleavage site in eukaryotic pre-mRNA splicing is determined by the overall 5’ splice region, not by the conserved 5’ GU. Cell 1987; 50: 237–246.

Anna A, Monika G. Splicing mutations in human genetic disorders: examples, detection, and confirmation. J Appl Genet 2018; 59: 253–268.

Aoyagi Y, Kobayashi H, Tanaka K, Ozawa T, Nitta H, Tsuji S. A de novo splice donor site mutation causes in-frame deletion of 14 amino acids in the proteolipid protein in Pelizaeus-Merzbacher disease. Ann Neurol 1999; 46: 112–115.

Baeza-Centurion P, Minana B, Schmiedel JM, Valcarcel J, Lehner B. Combinatorial genetics reveals a scaling law for the effects of mutations on splicing. Cell 2019; 176: 549–563 e523.

Bao S, Moakley DF, Zhang C. The splicing code goes deep. Cell 2019; 176: 414–416.

Bartolo C, Papp AC, Snyder PJ, Sedra MS, Burghes AH, Hall CD, Mendell JR, Prior TW. A novel splice site mutation in a Becker muscular dystrophy patient. J Med Genet 1996; 33: 324–327.

Blomen VA, Majek P, Jae LT, Bigenzahn JW, Nieuwenhuis J, Staring J, Sacco R, van Diemen FR, Olk N, Stukalov A, Marceau C, Janssen H, Carette JE, Bennett KL, Colinge J, Superti-Furga G, Brummelkamp TR. Gene essentiality and synthetic lethality in haploid human cells. Science 2015; 350: 1092–1096.

Burset M, Seledtsov IA, Solovyev VV. Analysis of canonical and non-canonical splice sites in mammalian genomes. Nucleic Acids Res 2000; 28: 4364–4375.

Cooper TA, Wan L, Dreyfuss G. RNA and disease. Cell 2009; 136: 777–793.

den Dunnen JT, Dalgleish R, Maglott DR, Hart RK, Greenblatt MS, McGowan-Jordan J, Roux AF, Smith T, Antonarakis SE, Taschner PE. HGVS Recommendations for the Description of Sequence Variants: 2016 Update. Hum Mutat 2016; 37: 564–569.

Den Uijl IE, Mauser Bunschoten EP, Roosendaal G, Schutgens RE, Biesma DH, Grobbee DE, Fischer K. Clinical severity of haemophilia A: does the classification of the 1950s still stand? Haemophilia 2011; 17: 849–853.

Erkelenz S, Theiss S, Kaisers W, Ptok J, Walotka L, Muller L, Hillebrand F, Brillen AL, Sladek M, Schaal H. Ranking noncanonical 5’ splice site usage by genome-wide RNA-seq analysis and splicing reporter assays. Genome Res 2018; 28: 1826–1840.

Findlay GM, Daza RM, Martin B, Zhang MD, Leith AP, Gasperini M, Janizek JD, Huang X, Starita LM, Shendure J. Accurate classification of *BRCA1* variants with saturation genome editing. Nature 2018; 562: 217–222.

Frischknecht H, Dutly F, Walker L, Nakamura-Garrett LM, Eng B, Waye JS. Three new beta-thalassemia mutations with varying degrees of severity. Hemoglobin 2009; 33: 220–225.

Fu XD, Ares M, Jr. Context-dependent control of alternative splicing by RNA-binding proteins. Nat Rev Genet 2014; 15: 689–701.

Goksuluk D, Korkmaz S, Zararsiz G, Karaagaoglu AE. easyROC: an interactive web-tool for ROC curve analysis using R language environment. The R Journal 2016; 8: 213–230.

Jaganathan K, Kyriazopoulou Panagiotopoulou S, McRae JF, Darbandi SF, Knowles D, Li YI, Kosmicki JA, Arbelaez J, Cui W, Schwartz GB, Chow ED, Kanterakis E, Gao H, Kia A, Batzoglou S, Sanders SJ, Farh KK. Predicting splicing from primary sequence with deep learning. Cell 2019; 176: 535–548 e524.

Kume K, Masamune A, Kikuta K, Shimosegawa T. [-215G>A; IVS3+2T>C] mutation in the *SPINK1* gene causes exon 3 skipping and loss of the trypsin binding site. Gut 2006; 55: 1214.

Lappalainen T, Scott AJ, Brandt M, Hall IM. Genomic analysis in the age of human genome sequencing. Cell 2019; 177: 70–84.

Lin JH, Masson E, Boulling A, Hayden M, Cooper DN, Férec C, Liao Z, Chen JM. 5’ splice site GC>GT variants differ from GT>GC variants in terms of their functionality and pathogenicity. bioRxiv 829010; doi: https://doi.org/10.1101/829010. 2019a.

Lin JH, Tang XY, Boulling A, Zou WB, Masson E, Fichou Y, Raud L, Le Tertre M, Deng SJ, Berlivet I, Ka C, Mort M, Hayden M, Leman R, Houdayer C, Le Gac G, Cooper DN, Li ZS, Férec C, Liao Z, Chen JM. First estimate of the scale of canonical 5’ splice site GT>GC variants capable of generating wild-type transcripts. Hum Mutat 2019b; 40: 1856–1873.

Matthews BW. Comparison of the predicted and observed secondary structure of T4 phage lysozyme. Biochim Biophys Acta 1975; 405: 442–451.

Mount SM, Avsec Z, Carmel L, Casadio R, Celik MH, Chen K, Cheng J, Cohen NE, Fairbrother WG, Fenesh T, Gagneur J, Gotea V, Holzer T, Lin CF, Martelli PL, Naito T, Nguyen TYD, Savojardo C, Unger R, Wang R, Yang Y, Zhao H. Assessing predictions of the impact of variants on splicing in CAGI5. Hum Mutat 2019; 40: 1215–1224.

Muller JS, Piko H, Schoser BG, Schlotter-Weigel B, Reilich P, Gurster S, Born C, Karcagi V, Pongratz D, Lochmuller H, Walter MC. Novel splice site mutation in the caveolin-3 gene leading to autosomal recessive limb girdle muscular dystrophy. Neuromuscul Disord 2006; 16: 432–436.

Parada GE, Munita R, Cerda CA, Gysling K. A comprehensive survey of non-canonical splice sites in the human transcriptome. Nucleic Acids Res 2014; 42: 10564–10578.

Pineda JMB, Bradley RK. Most human introns are recognized via multiple and tissue-specific branchpoints. Genes Dev 2018; 32: 577–591.

Ramalho AS, Beck S, Meyer M, Penque D, Cutting GR, Amaral MD. Five percent of normal cystic fibrosis transmembrane conductance regulator mRNA ameliorates the severity of pulmonary disease in cystic fibrosis. Am J Respir Cell Mol Biol 2002; 27: 619–627.

Raraigh KS, Han ST, Davis E, Evans TA, Pellicore MJ, McCague AF, Joynt AT, Lu Z, Atalar M, Sharma N, Sheridan MB, Sosnay PR, Cutting GR. Functional assays are essential for interpretation of missense variants associated with variable expressivity. Am J Hum Genet 2018; 102: 1062–1077.

Richards S, Aziz N, Bale S, Bick D, Das S, Gastier-Foster J, Grody WW, Hegde M, Lyon E, Spector E, Voelkerding K, Rehm HL, Committee ALQA. Standards and guidelines for the interpretation of sequence variants: a joint consensus recommendation of the American College of Medical Genetics and Genomics and the Association for Molecular Pathology. Genet Med 2015; 17: 405–424.

Scalet D, Maestri I, Branchini A, Bernardi F, Pinotti M, Balestra D. Disease-causing variants of the conserved +2T of 5’ splice sites can be rescued by engineered U1snRNAs. Hum Mutat 2019; 40: 48–52.

Scotti MM, Swanson MS. RNA mis-splicing in disease. Nat Rev Genet 2016; 17: 19–32.

Seyama K, Nonoyama S, Gangsaas I, Hollenbaugh D, Pabst HF, Aruffo A, Ochs HD. Mutations of the CD40 ligand gene and its effect on CD40 ligand expression in patients with X-linked hyper IgM syndrome. Blood 1998; 92: 2421–2434.

Shendure J, Findlay GM, Snyder MW. Genomic medicine-progress, pitfalls, and promise. Cell 2019; 177: 45–57.

Shi Y. Mechanistic insights into precursor messenger RNA splicing by the spliceosome. Nat Rev Mol Cell Biol 2017; 18: 655–670.

Starita LM, Ahituv N, Dunham MJ, Kitzman JO, Roth FP, Seelig G, Shendure J, Fowler DM. Variant interpretation: functional assays to the rescue. Am J Hum Genet 2017; 101: 315–325.

Stenson PD, Mort M, Ball EV, Evans K, Hayden M, Heywood S, Hussain M, Phillips AD, Cooper DN. The Human Gene Mutation Database: towards a comprehensive repository of inherited mutation data for medical research, genetic diagnosis and next-generation sequencing studies. Hum Genet 2017; 136: 665–677.

Vaz-Drago R, Custodio N, Carmo-Fonseca M. Deep intronic mutations and human disease. Hum Genet 2017; 136: 1093–1111.

Venet T, Masson E, Talbotec C, Billiemaz K, Touraine R, Gay C, Destombe S, Cooper DN, Patural H, Chen JM, Férec C. Severe infantile isolated exocrine pancreatic insufficiency caused by the complete functional loss of the *SPINK1* gene. Hum Mutat 2017; 38: 1660–1665.

Wang Z, Burge CB. Splicing regulation: from a parts list of regulatory elements to an integrated splicing code. RNA 2008; 14: 802–813.

Wu H, Boulling A, Cooper DN, Li ZS, Liao Z, Chen JM, Férec C. *In vitro* and *in silico* evidence against a significant effect of the *SPINK1* c.194G>A variant on pre-mRNA splicing. Gut 2017; 66: 2195–2196.

Zou WB, Boulling A, Masson E, Cooper DN, Liao Z, Li ZS, Ferec C, Chen JM. Clarifying the clinical relevance of SPINK1 intronic variants in chronic pancreatitis. Gut 2016; 65: 884–886.

