## Supplementary Table S1 for "The experimentally obtained functional impact assessments of GT>GC 5’ splice site variants differ markedly from those predicted"

**Supplementary Table S1.** Comparison of SpliceAI-predicted and experimentally demonstrated functional effects of the 45 disease-causing GT>GC (+2T>C) variants*

| Gene symbol | mRNA reference | Chr. | HG38 coordinate | Reference sequence | Variant | Generation of wild-type transcripts[a] | SpliceAI Delta score of donor loss | | |
|---|---|---|---|---|---|---|---|---|---|
| | | | | | | | +2T>C | +2T>A | +2T>G |
| *ABCC2* | NM_000392.5 | chr10 | 99811604 | T | c.1967+2T>C | No | 0.97 | 0.98 | 0.98 |
| *ACAT1* | NM_000019.3 | chr11 | 108146361 | T | c.1163+2T>C | No | 0.99 | 1 | 1 |
| *COQ8A* | NM_020247.4 | chr1 | 226984237 | T | c.1398+2T>C | No | 0.99 | 1 | 1 |
| *ALB* | NM_000477.6 | chr4 | 73417671 | T | c.1428+2T>C | No | 0.95 | 1 | 1 |
| *AR* | NM_000044.4 | chrX | 67711691 | T | c.2173+2T>C | No | 0.99 | 0.99 | 0.99 |
| *ART4* | NM_021071.3 | chr12 | 14842968 | A | c.144+2T>C | No | 0.97 | 0.99 | 0.99 |
| *ATP7A* | NM_000052.6 | chrX | 78011254 | T | c.1946+2T>C | No | 0.96 | 0.96 | 0.96 |
| *B9D1* | NM_015681.4 | chr17 | 19347782 | A | c.341+2T>C | No | 0.99 | 0.99 | 0.99 |
| *BTK* | NM_000061.2 | chrX | 101362171 | A | c.588+2T>C | No | 0.99 | 0.99 | 0.99 |
| *C8orf37* | NM_177965.3 | chr8 | 95269033 | A | c.155+2T>C | No | 0.92 | 0.92 | 0.92 |
| *CAV3* | NM_001234.4 | chr3 | 8733992 | T | c.114+2T>C | Yes (11%)[b] | 0.9 | 1 | 1 |
| *CD3E* | NM_000733.3 | chr11 | 118313876 | T | c.520+2T>C | Yes (1-5%)[c] | 0.99 | 0.99 | 0.99 |
| *CD40LG* | NM_000074.2 | chrX | 136654432 | T | c.346+2T>C | Yes (15%)[c] | 0.95 | 0.97 | 0.97 |
| *CLCN5* | NM_000084.4 | chrX | 50072590 | T | c.205+2T>C | No | 0.92 | 0.92 | 0.92 |
| *COL1A2* | NM_000089.3 | chr7 | 94426532 | T | c.3105+2T>C | No | 0.99 | 1 | 1 |
| *DGAT1* | NM_012079.5 | chr8 | 144318093 | A | c.751+2T>C | No | 0.99 | 0.99 | 0.99 |
| *DMD* | NM_004006.2 | chrX | 31657988 | A | c.8027+2T>C | Yes (10%)[c] | 0.63 | 0.99 | 0.99 |
| *DMD* | NM_004006.2 | chrX | 31206580 | A | c.9649+2T>C | No | 0.99 | 0.99 | 0.99 |
| *DNAL4* | NM_005740.2 | chr22 | 38780924 | A | c.153+2T>C | No | 0.98 | 0.99 | 0.99 |
| *GAA* | NM_000152.4 | chr17 | 80117111 | T | c.2331+2T>C | No | 0.94 | 0.97 | 0.97 |
| *HESX1* | NM_003865.2 | chr3 | 57198751 | A | c.357+2T>C[d] | No | 0.92 | 0.93 | 0.93 |
| *IL10RA* | NM_001558.3 | chr11 | 117994151 | T | c.688+2T>C | No | 0.96 | 0.97 | 0.97 |
| *ITGA8* | NM_003638.2 | chr10 | 15531048 | A | c.2982+2T>C | No | 0.95 | 0.99 | 0.99 |
| *IVD* | NM_002225.3 | chr15 | 40410799 | T | c.456+2T>C | No | 0.71 | 1 | 1 |
| *JAK3* | NM_000215.3 | chr19 | 17834569 | A | c.2350+2T>C | No | 0.71 | 0.97 | 0.99 |
| *LAMA2* | NM_000426.3 | chr6 | 129315952 | T | c.3924+2T>C | No | 0.97 | 0.97 | 0.97 |

| | | | | | | | | | |
|---|---|---|---|---|---|---|---|---|---|
| *LMAN1* | NM_005570.3 | chr18 | 59338758 | A | c.1149+2T>C | No | 0.97 | 0.97 | 0.97 |
| *MCM9* | NM_017696.2 | chr6 | 118827925 | A | c.1732+2T>C | No | 0.95 | 0.95 | 0.95 |
| *MMP2* | NM_004530.5 | chr16 | 55485429 | T | c.658+2T>C | No | 0.9 | 1 | 1 |
| *NCAPD2* | NM_014865.3 | chr12 | 6531078 | T | c.4120+2T>C | No | 0.88 | 1 | 1 |
| *NCF2* | NM_001127651.2 | chr1 | 183586893 | A | c.257+2T>C | No | 0.99 | 0.99 | 0.99 |
| *OPHN1* | NM_002547.2 | chrX | 68432865 | A | c.154+2T>C | No | 0.81 | 1 | 1 |
| *OTC* | NM_000531.5 | chrX | 38401430 | T | c.540+2T>C | No | 0.98 | 1 | 1 |
| *PCCB* | NM_000532.4 | chr3 | 136250560 | T | c.183+2T>C | No | 0.99 | 0.99 | 0.99 |
| *PCCB* | NM_000532.4 | chr3 | 136328859 | T | c.1498+2T>C | No | 1 | 1 | 1 |
| *PEX16* | NM_004813.2 | chr11 | 45910896 | A | c.952+2T>C | No | 0.97 | 0.99 | 0.99 |
| *PLP1* | NM_000533.4 | chrX | 103787968 | T | c.622+2T>C | No | 0.95 | 0.96 | 0.96 |
| *PLP1* | NM_000533.4 | chrX | 103788512 | T | c.696+2T>C | Yes (8%)[b] | 0.74 | 1 | 1 |
| *PNPLA2* | NM_020376.3 | chr11 | 823589 | T | c.757+2T>C | No | 0.95 | 0.99 | 0.99 |
| *PRDM5* | NM_018699.3 | chr4 | 120922514 | A | c.93+2T>C | No | 0.73 | 1 | 0.99 |
| *RFX6* | NM_173560.3 | chr6 | 116877954 | T | c.380+2T>C | No | 0.9 | 0.9 | 0.9 |
| *SCLT1* | NM_144643.3 | chr4 | 129039039 | A | c.290+2T>C | No | 0.95 | 0.95 | 0.95 |
| *SH2D1A* | NM_002351.4 | chrX | 124346781 | T | c.137+2T>C | No | 0.96 | 0.98 | 0.98 |
| *SLC26A2* | NM_000112.3 | chr5 | 149960981 | T | c.-26+2T>C | Yes (5%)[c] | 0.9 | 0.99 | 0.99 |
| *SPINK1* | NM_003122.3 | chr5 | 147828020 | A | c.194+2T>C[e] | Yes (10%)[b] | 0.35 | 0.99 | 1 |

*Refer to Lin et al. (2019b) for the original publications that reported the 45 variants.

[a]Relative expression level is indicated in parentheses.

[b]Expression level determined here by ImageJ using gel photos from the original publications.

[c]Expression level as described in the original publications.

[d]Identical to the *HESX1* IVS2+2T>C substitution in Supplementary Table S2.

[e]Identical to the *SPINK1* IVS3+2T>C substitution in Supplementary Table S2 and Table 2.
