## Supplementary Table S2 for "The experimentally obtained functional impact assessments of GT>GC 5’ splice site variants differ markedly from those predicted"

**Supplementary Table S2.** Comparison of SpliceAI-predicted and experimentally demonstrated functional effects of the 103 engineered GT>GC (+2T>C) substitutions

| Gene symbol | mRNA reference | Chr. | hg38 coordinate | Reference sequence | Variant[a] | Generation of wild-type transcripts[b] | SpliceAI Delta score of donor loss | | |
|---|---|---|---|---|---|---|---|---|---|
| | | | | | | | +2T>C | +2T>A | +2T>G |
| *ACTA1* | NM_001100.3 | chr1 | 229434002 | A | IVS1+2T>C | No | 0.84 | 0.99 | 0.99 |
| | | | 229432985 | A | IVS2+2T>C | No | 0.87 | 0.87 | 0.87 |
| | | | 229432554 | A | IVS3+2T>C | No | 0.26 | 1 | 0.96 |
| | | | 229432268 | A | IVS4+2T>C | No | 0.34 | 0.98 | 0.98 |
| | | | 229431719 | A | IVS6+2T>C | No | 0.28 | 1 | 1 |
| *AURKC* | NM_001015878.1 | chr19 | 57231789 | T | IVS2+2T>C | No | 0.92 | 0.96 | 0.96 |
| | | | 57232226 | T | IVS3+2T>C | No | 0.99 | 1 | 1 |
| | | | 57232682 | T | IVS4+2T>C | No | 0.99 | 0.99 | 0.99 |
| | | | 57233610 | T | IVS5+2T>C | No | 0.95 | 0.95 | 0.95 |
| | | | 57235060 | T | IVS6+2T>C | No | 0.8 | 0.8 | 0.8 |
| *CCDC103* | NM_213607.2 | chr17 | 44899861 | T | IVS1+2T>C | Yes | 0.82 | 0.82 | 0.82 |
| | | | 44901654 | T | IVS3+2T>C | No | 0.99 | 0.99 | 0.99 |
| *COX4I1* | NM_001861.4 | chr16 | 85799754 | T | IVS1+2T>C | No | 0.95 | 0.95 | 0.95 |
| | | | 85801280 | T | IVS2+2T>C | No | 0.92 | 0.93 | 0.93 |
| | | | 85805106 | T | IVS3+2T>C | No | 0.96 | 0.99 | 0.99 |
| | | | 85805866 | T | IVS4+2T>C | No | 0.82 | 1 | 1 |
| *COX7A1* | NM_001864.3 | chr19 | 36152391 | A | IVS1+2T>C | No | 0.95 | 0.96 | 0.96 |
| | | | 36151667 | A | IVS2+2T>C | No | 0.87 | 0.87 | 0.87 |
| *DBI* | NM_001079862.2 | chr2 | 119367062 | T | IVS1+2T>C | No | 0.99 | 0.99 | 0.99 |
| | | | 119368307 | T | IVS2+2T>C | Yes | 0.86 | 1 | 1 |
| | | | 119370804 | T | IVS3+2T>C | No | 0.87 | 1 | 1 |
| *DDIT3* | NM_004083.5 | chr12 | 57517704 | A | IVS2+2T>C | No | 0.98 | 1 | 1 |
| | | | 57517267 | A | IVS3+2T>C | No | 0.46 | 1 | 1 |
| *DNAJC19* | NM_145261.3 | chr3 | 180988176 | A | IVS2+2T>C | No | 0.94 | 0.95 | 0.95 |

| | | | | | | | | | |
|---|---|---|---|---|---|---|---|---|---|
| | | | 180985924 | A | IVS5+2T>C | Yes (42%) | 0.03 | 0.99 | 0.95 |
| *FABP1* | NM_001443.2 | chr2 | 88127949 | A | IVS1+2T>C | No | 0.96 | 0.99 | 0.99 |
| | | | 88126174 | A | IVS2+2T>C | No | 0.97 | 0.98 | 0.98 |
| | | | 88124492 | A | IVS3+2T>C | No | 0.98 | 0.98 | 0.98 |
| *FABP7* | NM_001446.4 | chr6 | 122779869 | T | IVS1+2T>C | No | 0.83 | 0.84 | 0.84 |
| | | | 122780465 | T | IVS2+2T>C | No | 0.64 | 1 | 1 |
| | | | 122781196 | T | IVS3+2T>C | No | 0.98 | 0.99 | 0.99 |
| *FATE1* | NM_033085.2 | chrX | 151716227 | T | IVS1+2T>C | Yes (84%) | 0.08 | 0.96 | 1 |
| | | | 151717401 | T | IVS2+2T>C | No | 0.98 | 0.99 | 0.99 |
| | | | 151721503 | T | IVS3+2T>C | No | 0.99 | 0.99 | 0.99 |
| | | | 151721983 | T | IVS4+2T>C | No | 0.92 | 1 | 1 |
| *FOLR3* | NM_000804.3 | chr11 | 72136122 | T | IVS2+2T>C | No | 0.96 | 1 | 1 |
| | | | 72139151 | T | IVS3+2T>C | No | 0.97 | 0.98 | 0.98 |
| | | | 72139484 | T | IVS4+2T>C | Yes | 0.45 | 1 | 1 |
| *HESX1* | NM_003865.2 | chr3 | 57199760 | A | IVS1+2T>C | Yes (2%) | 0.81 | 0.98 | 0.98 |
| | | | 57198751 | A | IVS2+2T>C[c] | No | 0.92 | 0.93 | 0.93 |
| | | | 57198389 | A | IVS3+2T>C | No | 0.96 | 1 | 1 |
| *HLA-DRA* | NM_019111.4 | chr6 | 32442695 | T | IVS2+2T>C | No | 0.99 | 0.99 | 0.99 |
| | | | 32443468 | T | IVS3+2T>C | No | 0.98 | 0.99 | 0.99 |
| | | | 32443923 | T | IVS4+2T>C | No | 0.93 | 0.99 | 0.99 |
| *IFI30* | NM_006332.4 | chr19 | 18175224 | T | IVS2+2T>C | No | 0.25 | 1 | 0.99 |
| | | | 18177294 | T | IVS5+2T>C | No | 0.98 | 1 | 1 |
| | | | 18177766 | T | IVS6+2T>C | No | 0.62 | 1 | 1 |
| *IFNL2* | NM_172138.1 | chr19 | 39268580 | T | IVS1+2T>C | No | 0.78 | 0.98 | 0.98 |
| | | | 39268860 | T | IVS2+2T>C | No | 0.88 | 0.97 | 0.96 |
| | | | 39269231 | T | IVS3+2T>C | No | 0.58 | 1 | 1 |
| | | | 39269639 | T | IVS4+2T>C | No | 0.99 | 0.99 | 0.99 |
| | | | 39269823 | T | IVS5+2T>C | Yes (5%) | 0.05 | 0.84 | 0.73 |
| *IL10* | NM_000572.3 | chr1 | 206772269 | A | IVS1+2T>C | No | 0.56 | 0.99 | 0.96 |

| | | | | | | | | | |
|---|---|---|---|---|---|---|---|---|---|
| | | | 206771354 | A | IVS2+2T>C | No | 0.23 | 0.99 | 1 |
| | | | 206770905 | A | IVS3+2T>C | Yes | 0.61 | 1 | 1 |
| | | | 206769827 | A | IVS4+2T>C | No | 0.44 | 1 | 1 |
| *LENG1* | NM_024316.2 | chr19 | 54159562 | A | IVS1+2T>C | No | 0.22 | 0.99 | 1 |
| | | | 54158280 | A | IVS2+2T>C | No | 0.89 | 1 | 1 |
| | | | 54156761 | A | IVS3+2T>C | No | 0.87 | 0.87 | 0.87 |
| *LY6G6F* | NM_001003693.1 | chr6 | 31707789 | T | IVS2+2T>C | No | 0.60 | 0.98 | 0.98 |
| | | | 31708136 | T | IVS3+2T>C | No | 0.81 | 0.81 | 0.81 |
| | | | 31710183 | T | IVS4+2T>C | No | 0.76 | 0.9 | 0.9 |
| | | | 31710420 | T | IVS5+2T>C | No | 0.3 | 0.3 | 0.3 |
| *MGP* | NM_000900.4 | chr12 | 14885729 | A | IVS1+2T>C | No | 0.98 | 0.99 | 0.99 |
| | | | 14884211 | A | IVS2+2T>C | Yes (80%) | 0.97 | 0.99 | 0.99 |
| *PIMREG* | NM_001195228.1 | chr17 | 6444490 | T | IVS1+2T>C | No | 0.97 | 0.99 | 0.99 |
| | | | 6445406 | T | IVS2+2T>C | No | 0.96 | 0.96 | 0.96 |
| | | | 6449409 | T | IVS4+2T>C | No | 0.99 | 1 | 1 |
| *PRSS2* | NM_002770.3 | chr7 | 142771024 | T | IVS1+2T>C | No | 0.81 | 0.99 | 0.98 |
| | | | 142772210 | T | IVS2+2T>C | No | 0.98 | 0.98 | 0.98 |
| | | | 142773521 | T | IVS3+2T>C | No | 0.75 | 1 | 0.99 |
| | | | 142774057 | T | IVS4+2T>C | No | 0.97 | 0.97 | 0.97 |
| *PSMC5* | NM_001199163.1 | chr17 | 63827925 | T | IVS1+2T>C | No | 0.93 | 0.98 | 0.98 |
| | | | 63830191 | T | IVS5+2T>C | No | 0.76 | 0.77 | 0.77 |
| | | | 63830503 | T | IVS6+2T>C | Yes (56%) | 0.31 | 0.98 | 1 |
| | | | 63830937 | T | IVS7+2T>C | No | 0.96 | 0.96 | 0.96 |
| | | | 63831228 | T | IVS8+2T>C | Yes (56%) | 0.21 | 1 | 1 |
| | | | 63831427 | T | IVS9+2T>C | No | 0.95 | 0.99 | 0.99 |
| | | | 63831618 | T | IVS10+2T>C | Yes (46%) | 0.83 | 1 | 1 |
| *RPL11* | NM_000975.5 | chr1 | 23691831 | T | IVS1+2T>C | No | 0.85 | 0.99 | 0.99 |
| | | | 23692761 | T | IVS2+2T>C | Yes | 0 | 0.87 | 0.86 |
| | | | 23693915 | T | IVS3+2T>C | Yes | 0.74 | 1 | 1 |

| | | | | | | | | | |
|---|---|---|---|---|---|---|---|---|---|
| | | | 23694793 | T | IVS4+2T>C | No | 0.94 | 0.94 | 0.94 |
| | | | 23695910 | T | IVS5+2T>C | No | 0.59 | 0.59 | 0.59 |
| *RPS17* | NM_001021.4 | chr15 | 82540424 | A | IVS1+2T>C | No | 0.95 | 0.98 | 0.98 |
| | | | 82539979 | A | IVS2+2T>C | No | 0.51 | 0.99 | 0.99 |
| | | | 82538878 | A | IVS3+2T>C | No | 0.99 | 1 | 1 |
| *RPS20* | NM_001023.3 | chr8 | 56074379 | A | IVS1+2T>C | No | 0.98 | 0.99 | 0.99 |
| | | | 56074058 | A | IVS2+2T>C | No | 0.95 | 0.99 | 0.99 |
| | | | 56073693 | A | IVS3+2T>C | No | 0.99 | 1 | 1 |
| *RPS27* | NM_001030.4 | chr1 | 153990804 | T | IVS1+2T>C | No | 0.93 | 0.94 | 0.94 |
| | | | 153991225 | T | IVS2+2T>C | Yes (63%) | 0.67 | 1 | 1 |
| | | | 153991678 | T | IVS3+2T>C | Yes | 0.98 | 1 | 1 |
| *SELENOS* | NM_203472.2 | chr15 | 101277340 | A | IVS1+2T>C | Yes | 0.81 | 1 | 1 |
| | | | 101276539 | A | IVS2+2T>C | No | 0.96 | 1 | 1 |
| | | | 101274590 | A | IVS4+2T>C | No | 0.97 | 0.98 | 0.98 |
| | | | 101274418 | A | IVS5+2T>C | Yes (14%) | 0.79 | 1 | 1 |
| | | | 101272762 | A | IVS6+2T>C | No | 0.64 | 0.64 | 0.64 |
| *SPINK1* | NM_003122.3 | chr5 | 147831521 | A | IVS1+2T>C | No | 0.85 | 0.86 | 0.86 |
| | | | 147829597 | A | IVS2+2T>C | No | 0.97 | 0.97 | 0.97 |
| | | | 147828020 | A | IVS3+2T>C[d] | Yes | 0.35 | 0.99 | 1 |
| *UQCRB* | NM_006294.4 | chr8 | 96235510 | A | IVS1+2T>C | No | 0.97 | 0.99 | 0.99 |
| | | | 96231772 | A | IVS3+2T>C | No | 1 | 1 | 1 |

[a]In accordance with the traditional IVS (InterVening Sequence; i.e., an intron) nomenclature as previously described (Lin et al. 2019b).

[b]Expression level (in parentheses), determined by quantitative RT-PCR analysis, was available for all +2T>C substitutions that generated only wild-type transcripts under the experimental conditions described in (Lin et al. 2019b).

[c]Identical to the *HESX1* c.357+2T>C variant in Supplementary Table S1.

[d]Identical to the *SPINK1* c.194+2T>C variant in Supplementary Table S1 and Table 1.
