## Supplementary Table S3 for "The experimentally obtained functional impact assessments of GT>GC 5’ splice site variants differ markedly from those predicted"

**Supplementary Table S3**. Confusion matrix using a threshold value of 0.85 derived from Supplementary Table S2

| | |
|---|---|
| **True positive (TP):** SpliceAI identifies a splicing event (>0.85) which produces no transcript | **False positive (FP):** SpliceAI identifies a splicing event (>0.85) which does produce transcript |
| **False negative (FN):** SpliceAI does not identify a splicing event (<0.85) which does not produce transcript | **True negative (TN):** SpliceAI does not identify a splicing event (<0.85) which does produce transcript |

| | |
|---|---|
| TP:57 | FP:3 |
| FN:27 | TN:16 |
