## Supplementary Fig. S1 for "The experimentally obtained functional impact assessments of GT>GC 5’ splice site variants differ markedly from those predicted"

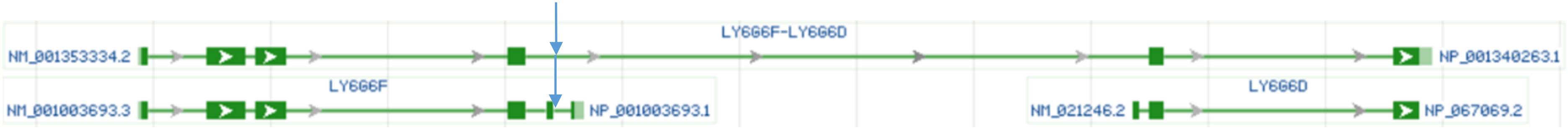


**Supplementary Figure S1.** Transcript isoforms of the *LY6G6F* gene. The approximate position of the discussed +2T site is indicated by arrow.
