## Supplementary Fig. S2 for "The experimentally obtained functional impact assessments of GT>GC 5’ splice site variants differ markedly from those predicted"

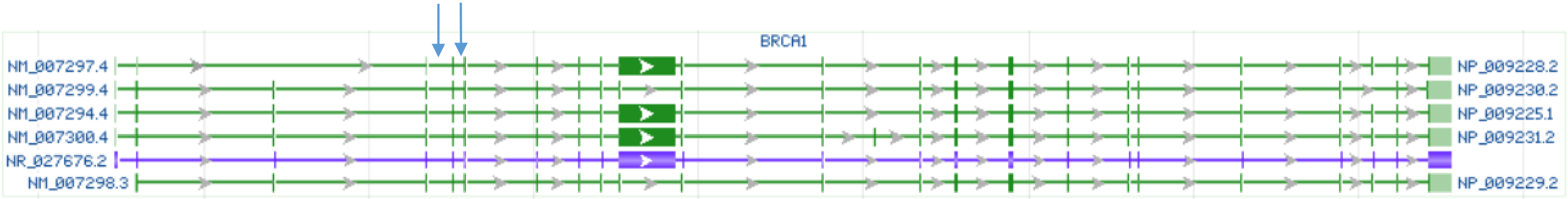


**Supplementary Figure S2.** Transcript isoforms of the *BRCA1* gene. The two discussed introns are indicated by arrows.
